# Bioactivation and detoxification of organophosphorus pesticides in freshwater planarians shares similarities with humans

**DOI:** 10.1101/2022.06.20.496885

**Authors:** Danielle Ireland, Christina Rabeler, TaiXi Gong, Eva-Maria S. Collins

## Abstract

Organophosphorus pesticides (OPs) are a chemically diverse class of insecticides that inhibit acetylcholinesterase (AChE). Many OPs require bioactivation to their active oxon form via cytochrome P450 to effectively inhibit AChE. OP toxicity can be mitigated by detoxification reactions performed by carboxylesterase and paraoxonase. The relative extent of bioactivation to detoxification varies among individuals and between species, leading to differential susceptibility to OP toxicity. Because of these species differences, it is imperative to characterize OP metabolism in model systems used to assess OP toxicity to adequately evaluate potential human hazard. We have shown that the asexual freshwater planarian *Dugesia japonica* is a suitable model to assess OP neurotoxicity and developmental neurotoxicity via rapid, automated testing of adult and developing organisms in parallel using morphological and behavioral endpoints. *D. japonica* has two cholinesterase enzymes with intermediate properties between AChE and butyrylcholinesterase that are sensitive to OP inhibition. Here, we demonstrate that this planarian contains the major OP metabolic machinery to be a relevant model for OP neurotoxicity studies. Adult and regenerating *D. japonica* can bioactivate chlorpyrifos and diazinon into their respective oxons. Significant AChE inhibition was only observed after *in vivo* metabolic activation but not when the parent OPs were directly added to planarian homogenate. Additionally, we found that *D. japonica* has both carboxylesterase and paraoxonase activity. Using specific chemical inhibitors, we show that carboxylesterase activity is distinct from cholinesterase activity. Taken together, these results further support the use of *D. japonica* for OP toxicity studies.

## Introduction

Organophosphorus pesticides (OPs) are inexpensive and readily-available pesticides, and are important tools for fighting vector-borne diseases, protecting crops from pests and diseases, and maintaining public parks and other urban areas (Nicolopoulou-Stamati et al. 2016). Hence, they are present in both agricultural and domestic domains (EUROSTAT 2016; Atwood and Paisley-Jones 2017). This class of compounds is defined by its inhibition of acetylcholinesterase (AChE, EC 3.1.1.7), an enzyme that breaks down the neurotransmitter acetylcholine (ACh) and is integral to cognitive, peripheral autonomic, and somatic motor function (Reiner and Radić 2000). In insects and in humans, inhibition of AChE leads to cholinergic toxicity, which ultimately results in paralysis and death (Russom et al. 2014). Each year, an estimated 3 million people are exposed to OPs, including 300,000 attributable fatalities (Robb and Baker 2021). Additionally, a growing number of studies have suggested that chronic, low-dose exposure to these pesticides correlate with adverse neurodevelopmental and neurodegenerative outcomes (Rauh et al. 2011; Muñoz-Quezada et al. 2013; Shelton et al. 2014; González-Alzaga et al. 2014; Sánchez-Santed et al. 2016; Burke et al. 2017; Sagiv et al. 2021). Despite the global usage of OPs, there is a lack of understanding of the health consequences of low-dose OP exposure because traditional toxicity testing is extremely slow and expensive.

Therefore, there has been a push to reduce, replace, and refine (3 R’s) the use of mammalian testing using methodologies that are inexpensive and higher throughput. New approach methods (NAMs), including *in vitro, in silico*, and non-vertebrate *in vivo* models (also referred to as “non-animal” organismal models), are being assessed as part of integrated testing strategies, with the goal to develop a robust, rapid test battery that is on par with or exceeds the relevancy of current mammalian models to human toxicity (Rovida et al. 2015; Sachana et al. 2021). The asexual freshwater planarian *Dugesia japonica* is an organismal invertebrate model that is uniquely suited to neurotoxicity and developmental neurotoxicity studies (Hagstrom et al. 2015, 2016; Zhang et al. 2019a; Ireland et al. 2020). This planarian reproduces exclusively asexually via binary fission into a head and tail piece. The resulting offspring regenerate the missing body parts, including a functional brain in the tail piece (Cebrià 2007; Ross et al. 2017). Thus, neuroregeneration in planarian tails serves as a rapid and fundamentally analogous process to neurodevelopment in vertebrates (Cebrià et al. 2002; Cebrià 2007; Hagstrom et al. 2016; Ross et al. 2017) and can be induced at will using amputation. Because of the similar size of intact and decapitated planarians, adult and regenerating planarians can be assessed in parallel with the same assays to identify effects that are specific to neurodevelopment (Hagstrom et al. 2015; Zhang et al. 2019b, a). Using an automated screening platform assessing multiple morphological and behavioral outcomes, we have shown that *D. japonica* are sensitive to OP exposure (Hagstrom et al. 2015; Zhang et al. 2019a) and that exposure to the OP chlorpyrifos (CPF) causes different effects in regenerating planarians than in adult planarians (Hagstrom et al. 2015; Zhang et al. 2019a).

The toxicity of OPs is regulated in large part by the extent to which they are metabolized. Many OPs, including CPF, require desulfuration (bioactivation) by cytochrome P450 (CYP) into their active oxon form to be able to effectively inhibit AChE. For example, the affinity of CPF-oxon (CPFO) to recombinant human AChE is three times greater than that of CPF (Amitai et al. 1998). Several human CYPs are responsible for the desulfuration of a single OP into its oxon, with varying CYPs activating different OPs. The relative importance of specific CYPs for OP bioactivation can depend on OP concentration (Buratti et al. 2003). Additionally, OPs can be detoxified by CYPs, paraoxonase-1 (PON1), and carboxylesterase (CES) among others (Alejo-González et al. 2018). Humans have three paraoxonase genes, of which PON1, a calcium-dependent aryldialkylphosphatase (EC 3.1.8.1), is the most important for OP detoxification (Taler-Verčič et al. 2020). PON1 is named for its ability to detoxify paraoxon, the active metabolite of the OP parathion (Aldridge 1953; Mackness et al. 1998). PON1 activity has been shown to be a major determinant of OP toxicity, including in humans (Costa et al. 1999). Species differences in PON1 activity exist, which have also been shown to be substrate specific (Carr et al. 2015). In addition to its role in xenobiotic metabolism, PON1 decreases oxidation of low-density lipoproteins and has thus been suggested to play a protective role in several human diseases, including diabetes mellitus and atherosclerosis (Shunmoogam et al. 2018).

CES (EC 3.1.1.1) is a serine hydrolase, like AChE, and hydrolyzes various carboxylic esters, including endogenous esters in lipid and glucose metabolism and xenobiotics (Hatfield et al. 2016; Wang et al. 2018). CES inhibits the function of some OPs, such as malathion and isocarbophos, by destructively phosphorylating their active sites (Buratti and Testai 2005; Zhuang et al. 2014). CES is also susceptible to OP inhibition by compounds like CPFO, paraoxon, and methyl paraoxon (Crow et al. 2008; Hatfield and Potter 2011). CES sequesters these pesticides from other targets such as AChE (Maxwell and Brecht 2001), thus, acting as a bioscavenger. This duality is important to understand when evaluating toxicity of mixtures of OPs – as is commonly the case in real-life exposures. For example, co-administration of CPF and malathion leads to enhanced toxicity compared to malathion alone, because CPF inhibits CES from detoxifying malathion (Jansen et al. 2009).

The relative extent of bioactivation to detoxification varies greatly between species and impacts a species’ relative sensitivity to OPs. For example, acephate is metabolized to its more toxic metabolite methamidophos in both insects and in mammals. However, in mammals, methamidophos triggers a negative feedback loop inhibiting further bioactivation, leading to reduced acephate toxicity compared to insects (Mahmoud Mahajna et al. 1997). Moreover, some strains of insects have gained OP-resistance which has been directly linked to changes in CES activity (Sun et al. 2005; Cui et al. 2007; Gong et al. 2017). Thus, it is imperative to characterize OP metabolism in model systems used to assess OP toxicity to be able to contextualize results and predict potential human toxicity.

While planarians have been used for over a century for pharmacological and toxicological studies (Best et al. 1981; Best and Morita 1991; Hagstrom et al. 2016), little is known about how they metabolize xenobiotics and whether they have the relevant targets of OP toxicity. We have previously shown that *D. japonica* has two cholinesterase enzymes (DjChE-1 and DjChE-2) which are functionally similar to both AChE and butylrylcholinesterase (BChE) (Hagstrom et al. 2017, 2018). These enzymes show similar OP inhibition and reactivation kinetics to mammalian AChE and are evolutionary ancestors of mammalian AChE and BChE (Hagstrom et al. 2017). Correlations between some phenotypic outcomes and decreased ChE activity have been shown after *in vivo* exposure to the carbamate physostigmine, diazinon (DZN) and some organophosphorus flame retardants (Hagstrom et al. 2018; Zhang et al. 2019b). Since DZN does require bioactivation into its oxon form, diazoxon (DZNO) (Poet et al. 2003), inhibition in these conditions suggests that planarians can bioactivate OPs. To further verify the use of *D. japonica* as a model to study OP toxicity, we tested whether *D. japonica* planarians contain the main metabolic machinery necessary to bioactivate and detoxify OPs using a series of colorimetric assays to detect and differentiate cholinesterase, PON1 and CES activity. We find that *D. japonica* can bioactivate CPF and DZN at all developmental stages, shows PON1 activity and contains CES activity that is distinct from the previously characterized cholinesterase activity. Taken together, these results validate the use of *D. japonica* for OP toxicity studies.

## Materials and Methods

### Planarian culture

Freshwater asexual planarians of the species *Dugesia japonica* were used for all experiments. Planarians were maintained in 1x Instant Ocean (IO, Blacksburg, VA) in Tupperware containers at 20°C in a Panasonic refrigerated incubator in the dark. Animals were fed organic chicken or beef liver 1-2x/week and cleaned twice a week when not used for experiments. Animals were starved for at least 5 days before experiments.

### Cholinesterase activity assays

To determine whether *D. japonica* planarians could bioactivate CPF or DZN into their active oxon forms, we compared DjChE levels after exposure to 1 mM CPF or 1 µM CPFO, and to 31.6 µM DZN or 0.1 µM DZNO (Table 1). These OPs/oxons were chosen because of the known efficacy of the respective oxons to inhibit DjChE at these concentrations (Hagstrom et al. 2017). Stocks were prepared in 100% dimethyl sulfoxide (DMSO, Sigma-Aldrich, Saint Louis, MO). The final DMSO concentration in the exposure solution was ≤ 0.5%. To quantify *in vivo* inhibition of ChE activity, 30 similarly-sized planarians were exposed to the respective inhibitor for 30-55 minutes. For the exposure, planarians were kept in 12-well plates (Genesee Scientific, San Diego, CA), with 6 planarians per well and a total volume of 1.2 mL. At the end of exposure, the planarians were washed three times with IO water and homogenized in 1% Triton X-100 (Sigma-Aldrich) in PBS as previously described (Hagstrom et al. 2017, 2018). Homogenates were kept on ice until activity measurements were made. To quantify *in vitro* inhibition of ChE activity, a wild-type planarian homogenate was prepared as previously described (Hagstrom et al. 2017). Then, 10X stock of the respective inhibitor was added to the homogenate to achieve the final 1X concentration. After gently vortexing, the inhibitor/homogenate mix was incubated for 40 minutes at room temperature and then the ChE activity was immediately quantified. Forty minutes was used to be roughly equivalent to the 30 minute *in vivo* exposure + 10 minutes needed for homogenization. These experiments were performed on either intact/adult planarians or regenerating planarians which had been amputated pre-pharyngeally within 3 hours of exposure. Levels of acetylthiocholine (ATCh) catalysis (ChE activity) were determined either as previously described (Hagstrom et al. 2017) or using an Acetylcholinesterase Activity Assay Kit (Sigma-Aldrich). Absorbance was read at 412 nm every minute for 10 minutes using a VersaMax (Molecular Devices, San Jose, CA) spectrophotometer. ChE activity was calculated as the rate of change of absorbance per minute during the linear portion of the reaction. ChE activity was normalized by protein concentration as determined by a Coomassie (Bradford) protein assay kit (Thermo Scientific, Waltham, MA) and compared to vehicle-exposed samples (set at 100% activity). Activity measurements were performed with at least 3 technical replicates per condition and at least 2 independent experiments (biological replicates).

**Table 1.** Inhibitors used in this study.

| Chemical | CAS | Concentration | Class | Supplier | Purity (%) |
| --- | --- | --- | --- | --- | --- |
| Chlorpyrifos (CPF) | 2921-88-2 | 1 mM | OP | Sigma-Aldrich | 100 |
| Chlorpyrifos oxon (CPFO) | 5598-15-2 | 1 μM | Active metabolite | Chem Service (West Chester, PA) | 98 |
| Diazinon (DZN) | 333-41-5 | 31.6 μM | OP | Sigma-Aldrich | 98 |
| Diazoxon (DZNO) | | 0.1 μM | Active metabolite | Chem Service | 97 |
| Methyl paraoxon | 950-35-6 | 10 mM<br>2 mM | B-esterase inhibitor<br>PTEH substrate | Sigma-Aldrich | 100 |
| Benzil | 134-81-6 | 1.6 mM | Carboxylesterase inhibitor | Fisher Scientific (Hampton, NH) | 99 |
| Tacrine hydrochloride | 1684-40-8 | 3.13 mM | Cholinesterase inhibitor | Cayman Chemical (Ann Arbor, MI) | ≥98 |
| Ethylenediamine tetraacetic acid (EDTA) | 60-00-4 | 1 μM | Calcium chelator | Sigma-Aldrich | 99 |

To visualize ChE activity during the course of regeneration, intact planarians were amputated pre-pharyngeally on day 1. Four planarians were individually fixed and stained for qualitative detection of ATCh catalysis (DjChE activity) on each subsequent day of regeneration (day 2-7) as previously described (Hagstrom et al. 2017, 2018). Staining was performed for 6 hours to ensure a robust signal. Samples were imaged in brightfield using a Point Grey Flea®3 color camera (Teledyne FLIR, Wilsonville, OR) mounted on a Leica S8 APO stereo microscope (Leica Camera, Wetzlar, Germany). Representative images are shown.

### Paraoxonase-1 (PON1) activity assays

PON1 activity was determined *in vitro* using an adapted version of the phosphoric triester hydrolase (PTEH) assay described in (Anspaugh and Roe 2002). Wild-type intact planarians were homogenized as described above but in cold 100 mM Tris-Cl, pH 8. Dilutions of the homogenate were performed in Tris buffer. The reaction was performed in 48-well plates and was initiated by combining 289 µl of homogenate with 11 µl 2 mM methyl paraoxon. After a 45-minute incubation at room temperature with rotation, the absorbance at 405 nM was measured using a VersaMax plate reader. The absorbance of the samples was compared to a standard curve created using different concentrations of p-nitrophenol. Protein concentration was determined by a Coomassie (Bradford) protein assay kit. PTEH specific activity was calculated as 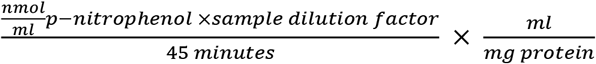. To determine the specificity of the reaction, 2 µl of 0.5 mM EDTA was pre-incubated with 1 ml of the homogenate for 10 minutes before the PTEH activity. Since PON1 requires calcium for its enzymatic activity (Anspaugh and Roe 2002), EDTA was used as a negative control to inhibit PTEH activity.

### Carboxylesterase (CES) activity assays

*In vitro* levels of CES activity were quantified using an adapted version of the carboxylic ester hydrolase (CEH) assay from (Anspaugh and Roe 2002), which measures general B-esterase activity, including both ChE and CES. Wild-type planarian homogenates were prepared in the same way as in the ChE activity assays (Hagstrom et al. 2017). Clarified homogenate (70 µl) was pre-incubated with either 5 µl 100 mM sodium phosphate buffer or 10 mM methyl-paraoxon (an irreversible inhibitor of B-type esterases) (Aldridge 1953; Montella et al. 2012). After a 10-minute incubation with rotation, 100 µl of 1 mM 1-napthyl acetate was then added to the samples, which were subsequently incubated for 15 minutes. When hydrolyzed by CES, 1-naphthyl-acetate is converted to 1-naphthol, whose bluish color is detectable at 595 nm. The samples and 1-naphthol standards were incubated with 25 µl 0.3% Fast Blue B Salt (Sigma-Aldrich), containing 3.4% sodium dodecyl sulfate (Life Technologies, Gaithersburg, MD) for 10 minutes to allow color formation. Absorbance was then immediately measured at 595 nm using a VersaMax plate reader and compared to the 1-naphthol standard curve. Protein concentration was determined by a Coomassie (Bradford) protein assay kit. CEH activity (nmol/(min X mg protein)) was calculated as 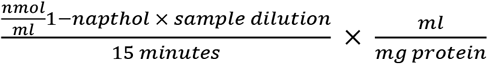. All experiments were conducted at room temperature. All standards and sample activity measurements were performed with 3 technical replicates per condition and in 2 independent experiments.

To distinguish ChE and CES activity, specific inhibitors of either ChE or CES activity were pre-incubated with the planarian homogenate for 10 minutes (instead of methyl paraoxon mentioned above), followed by an Ellman or CEH assay as described above. The CES-specific inhibitor benzil (Wang et al. 2018) was used to identify ChE activity, whereas tacrine, a ChE-specific inhibitor (Grishchenko et al. 2022), was used to measure CES-specific activity. 16 mM benzil (10X stock) was dissolved in 100% ethanol. Tacrine was prepared at 31.25 mM (10X stock) in 1% Triton X-100 in PBS. 10X stocks were added to homogenate to achieve the desired final 1X concentrations. Concentrations of tacrine were determined based on previous inhibition experiments with DjChE (Hagstrom et al. 2017). Inhibition was calculated by comparing to the activity from the homogenates exposed to the respective solvent (control), which were set as 100% activity. 10% ethanol was found to have no effect on either CEH or AChE activity alone.

### Analysis

Analysis of the colorimetric data was performed in Excel. Plots were created in MATLAB (MathWorks, Natick, MA). Artwork and figures were made in Inkscape (Free Software Foundation, Inc., Boston, MA).

## Results

### ChE activity during D. japonica regeneration

Planarians provide the unique ability to study both adult and regenerating/developing organisms in parallel, allowing for differentiation of developmental-specific effects (Hagstrom et al. 2015; Zhang et al. 2019a). While the expression and activity of DjChE has been studied in adult planarians (Hagstrom et al. 2018), how DjChE activity changes over the course of regeneration is unknown. Thus, we amputated *D. japonica* planarians on day 1 and qualitatively visualized DjChE activity in the regenerating tail pieces every day for the next 7 days (**Fig. 1a**). DjChE activity was retained in the ventral nerve cords in the tail piece throughout regeneration. DjChE activity could be seen in the regenerating brain starting at day 4 and gradually increased over the next 7 days. These results were verified quantitatively using Ellman assays (Ellman et al. 1961) (**Fig. 1b**). When compared to adult controls, DjChE activity dips immediately after amputation due to loss of the head, in which the highest density of DjChE is found (Hagstrom et al. 2018). DjChE activity then steadily increases over the course of regeneration, plateauing after about 1 week.

**Fig. 1.**
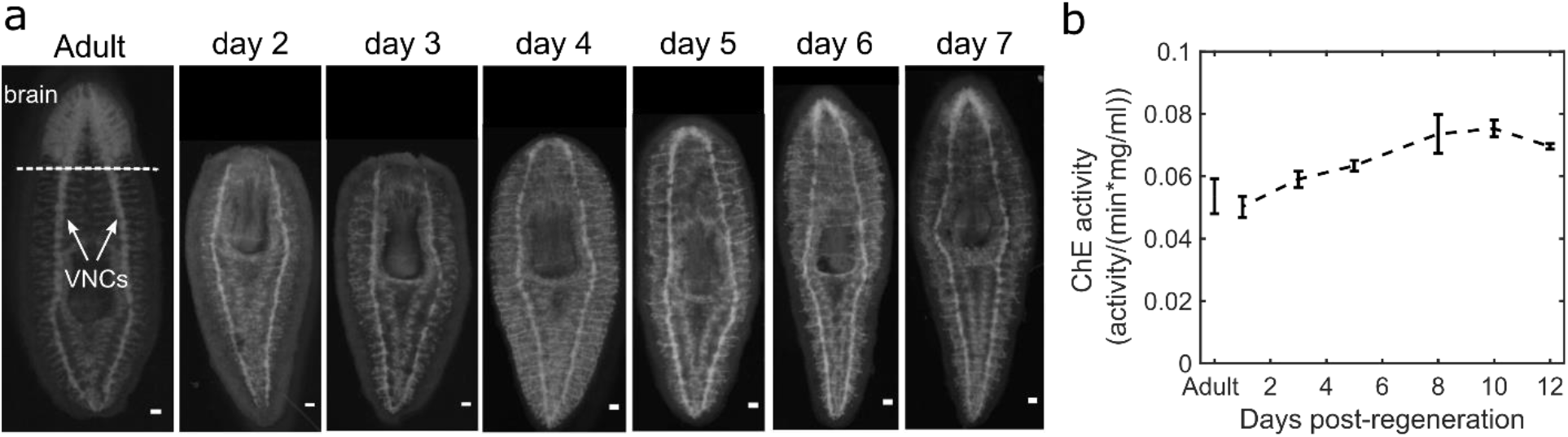
DjChE activity during head regeneration. a) Adult planarians were amputated on day 1. Dotted line indicates amputation plane. Arrows point to ventral nerve cords (VNCs). DjChE activity was visualized at each day of regeneration for the first 7 days. Scale bars: 100 µm. b) Adult planarians were amputated on day 1. Ellman assays were performed on regenerating planarians at different points during regeneration. DjChE activity was normalized by protein activity. Error bars indicate standard deviation of 3 technical replicates

### D. japonica can bioactivate OPs into their oxons at all developmental stages

To test whether OPs are bioactivated in *D. japonica*, we performed Ellman assays (Ellman et al. 1961) to quantify the catalytic activity of DjChE in the presence of parent OPs (CPF and DZN) or their active oxons (CPFO and DZNO, respectively). We have previously shown that these oxons are potent inhibitors of DjChE *in vitro* (Hagstrom et al. 2017). When the inhibitors were added directly to planarian homogenates (*in vitro* exposure), DjChE activity was strongly inhibited by the oxons but not by the parent OPs (**Fig. 2**), demonstrating that CPF and DZN do not significantly inhibit DjChE activity directly. As integral membrane proteins (Szczesna-Skorupa and Kemper 2008), CYPs are likely not active in the crude planarian homogenates which contains mostly soluble proteins. In contrast, if planarians were pre-incubated for 30-55 minutes with these concentrations and then homogenized (*in vivo* exposure) and used in the Ellman assay, both the parent and oxon forms inhibited DjChE activity (**Fig. 2**), demonstrating that bioactivation of CPF into CPFO and DZN into DZNO occurs in *D. japonica*. Notably, these trends were the same whether we used intact/adult planarians (**Fig. 2a)** or regenerating planarians (**Fig. 2b)**, demonstrating that bioactivation occurs at all developmental stages in *D. japonica*.

**Fig. 2.**
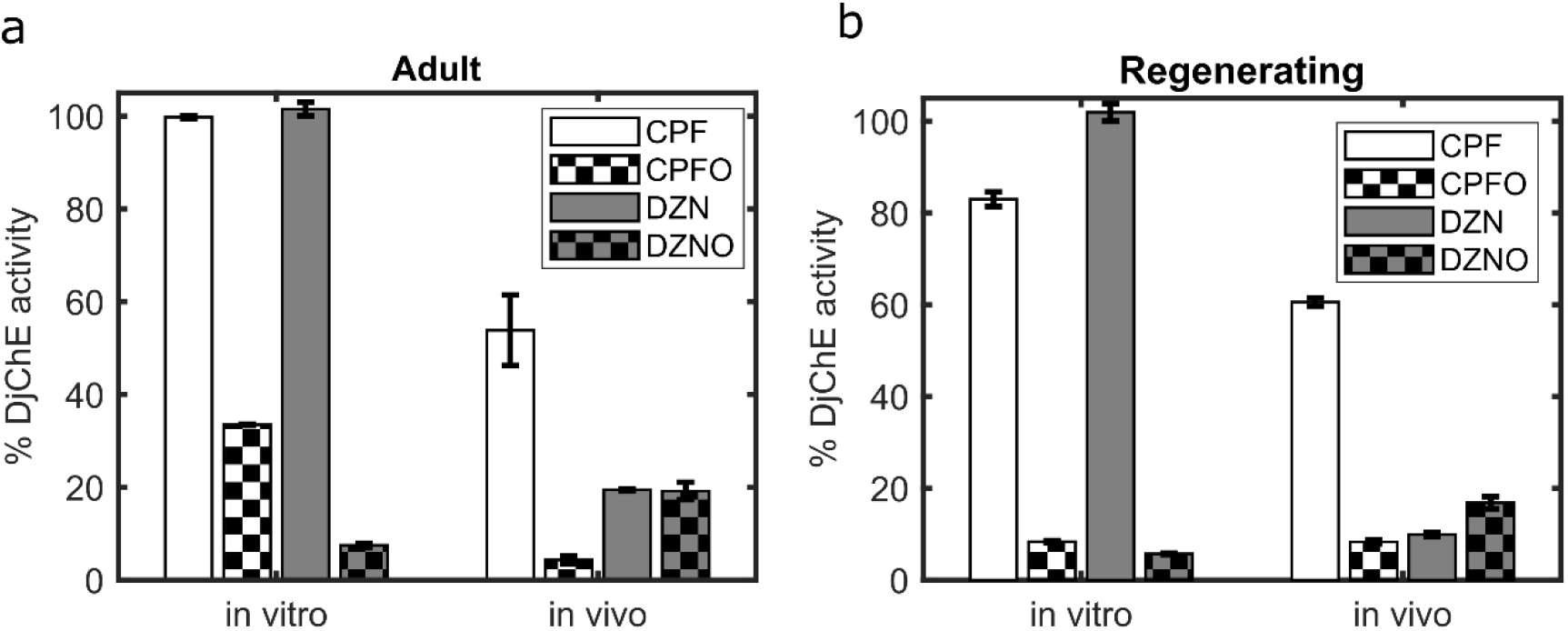
Bioactivation of CPF and DZN at all developmental stages in *D. japonica*. a-b) CPF and DZN do not inhibit DjChE directly *in vitro* but do *in vivo* in (a) adult and (b) regenerating planarians. Error bars are standard error of at least 6 replicates over at least 2 experiments (biological replicates)

### D. japonica has PON1 activity

To test whether *D. japonica* have PON1 activity to break down paraoxon, we performed a colorimetric PTEH assay (Anspaugh and Roe 2002). *D. japonica* had 60 ± 7 (mean ± standard error) pmol/(min*mg protein) PTEH specific activity (**Fig. 3**). The calcium chelator EDTA was used to test the specificity of the activity since PON1 requires calcium as a cofactor for its enzymatic activity (Anspaugh and Roe 2002). *D. japonica* PTEH activity was effectively inhibited (76% reduction in activity) by EDTA.

**Fig. 3.**
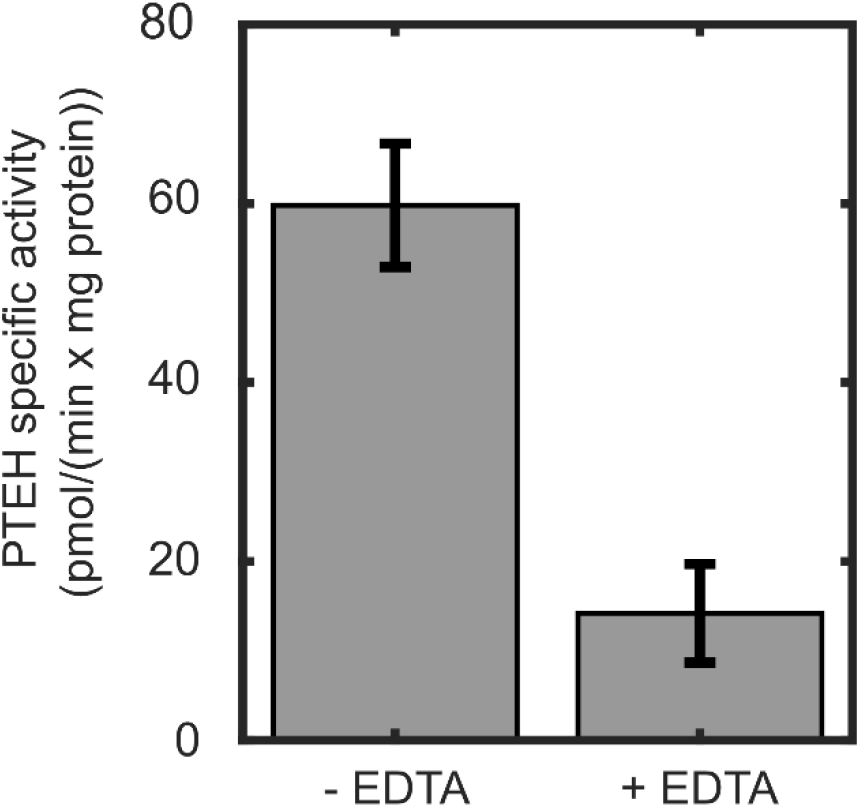
*D. japonica has PON1 activity*. Quantification of PTEH activity (Anspaugh and Roe 2002) from adult wild-type *D. japonica* planarians. The specificity of the activity was determined by the addition of 1 µM EDTA to inhibit the calcium-dependent activity of PON1. Errors bars indicate standard error of 6 technical replicates over 2 independent experiments (biological replicates)

### D. japonica has CES activity that is distinct from DjChE activity

We performed a colorimetric CEH assay (Anspaugh and Roe 2002) to determine whether *D. japonica* contains CES activity. We found that *D. japonica* have approximately 24 ± 1.7 (mean ± standard error) nmol(min*mg protein) CEH activity (**Fig. 4A**). CEH activity can be performed by a large array of enzymes, in which B-esterases (including CES and ChE) are a subset (Anspaugh and Roe 2002). We found that planarian CEH activity was strongly inhibited (96% compared to the untreated control) by methyl paraoxon, a specific inhibitor of B-esterases (Aldridge 1953; Anspaugh and Roe 2002; Montella et al. 2012), suggesting that this activity is performed primarily by B-esterases.

**Fig. 4.**
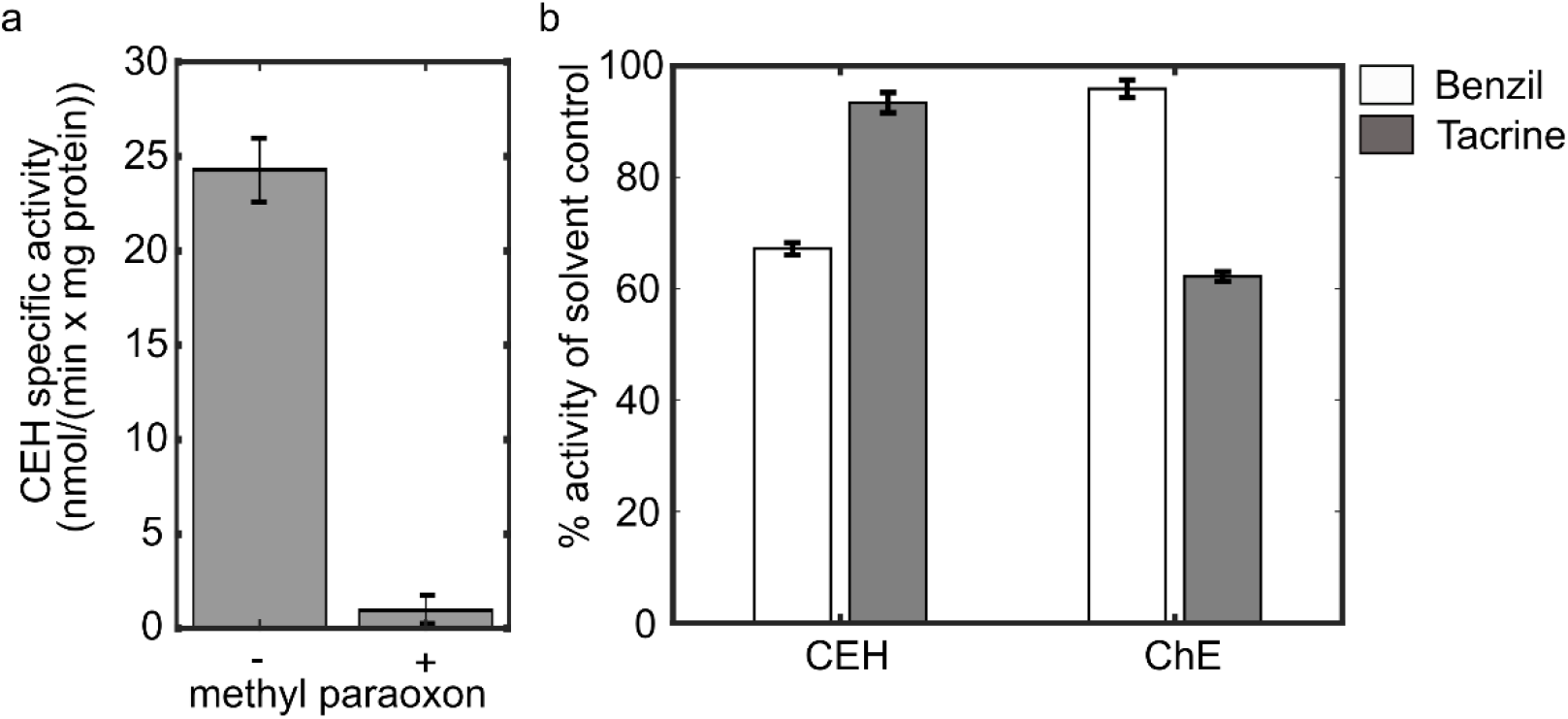
*D. japonica* has CES activity that is distinct from ChE activity. a) Quantification of CEH activity (Anspaugh and Roe 2002) from homogenates of wild-type adult *D. japonica*. CEH activity was inhibited by 10 mM methyl paraoxon. b) 1.6 mM benzil inhibits CEH activity but not ChE activity, whereas 3.13 mM tacrine inhibits ChE activity but not CEH activity, suggesting the two enzymatic activities are distinct. Errors bars indicate standard error of 6 technical replicates over 2 independent experiments (biological replicates)

Although 1-naphthyl acetate is sometimes considered a specific substrate for CES, it can also be hydrolyzed by AChE (Chowdhary et al. 2018). Thus, to determine if the detected CEH activity was distinct from ChE activity, we compared the catalysis of 1-naphthyl acetate and ATCh in the presence of a CES-specific inhibitor benzil (Wang et al. 2018) or a ChE-specific inhibitor tacrine (**Fig. 4B**). We have previously shown that DjChE activity is sensitive to tacrine inhibition (Hagstrom et al. 2017). The CES-specific inhibitor benzil (1.6 mM) inhibited CEH activity by ∼33% while causing very minor inhibition (4%) of ChE activity in the Ellman assay. Conversely, 3.13 mM tacrine caused inhibition of DjChE activity in the Ellman assay (38%) but negligible (7%) inhibition in the CEH assay. This differential action of the two inhibitors suggests that some of the CEH activity is due to CES and that CES activity is distinct from ChE activity in *D. japonica*.

## Discussion

### *D. japonica* can metabolize CPF and DZN at all life stages

Chemical toxicity depends on metabolism. For the OPs studied here, CPF and DZN, the major routes of metabolism are comprised of bioactivation via CYP enzymes into their respective oxon forms (CPFO and DZNO) which inhibit AChE, and detoxification into their primary metabolites (3,5,6-trichloro-2-pyridinol (TCP) for CPF and 2-isopropyl-4-methyl-6-hydroxypyrimidine (IMHP) for DZN), via CYPs, CES, and PON1 (Poet et al. 2003). In humans, CYP2B6 was found to have the highest activity for desulfuration of CFP and CYP1A1 for DZN, whereas CYP2C19 is the primary CYP involved in detoxification for both OPs (Tang et al. 2001; Ellison et al. 2012).

OP toxicity is augmented or diminished by the relative balance of bioactivation to detoxification. Because of differences in gene expression and activity of these enzymes, some parts of the human population (e.g., children or specific ethnic groups) are more vulnerable to OP exposure than others (Tang et al. 2001; Costa et al. 2013; Gómez-Martín et al. 2015; Wang et al. 2018). Gender differences in enzyme activity and thus OP susceptibility also exist (Tang et al. 2001). Similarly, polymorphisms in the genes encoding CES have been shown to underly OP resistance in certain strains of insects, including mosquitoes and cotton aphids (Sun et al. 2005; Cui et al. 2007; Gong et al. 2017). Interspecies differences in enzyme activity are even greater than intraspecies variability, and thus different species can exhibit vastly disparate susceptibility when exposed to certain OPs (Maxwell et al. 1987; Veronesi and Ehrich 1993). For example, mice have been shown to have dramatically reduced sensitivity to OP nerve agents due to the presence of plasma CES, which is not found in humans. Because of this, CES knockout mice are used as a more appropriate model of OP toxicity (Duysen et al. 2011). Similarly, inhibition of CES activity has been shown to negate species differences in soman-induced lethality across several rodent species (Maxwell et al. 1987). Moreover, species with low PON1 activity, such as birds, have increased sensitivity to OPs (Costa et al. 2008). The relative importance of these different enzymes is dependent on the OP as not all OPs are metabolized in the same manner. For example, CES has been suggested to be more relevant for protection against organophosphorus nerve agents than OPs (Lockridge et al. 2016). Because of these known differences, it is imperative to understand the extent that xenobiotic metabolism occurs in model systems used for toxicity screening of different chemical domains, so that results can be translated to human health.

Here, we showed that the freshwater planarian *D. japonica* has the metabolic machinery to bioactivate and detoxify OPs. CYP enzymes are integral membrane proteins localized to the endoplasmic reticulum (Szczesna-Skorupa and Kemper 2008) and thus are not active in most crude homogenate preparations, as used here, which mainly consist of soluble proteins, including most esterases. CPF and DZN could not inhibit DjChE directly in *in vitro* homogenates but could after at least 30-minute *in vivo* incubation in the whole organism. Because of the similar timing used in the two experiments, the *in vivo* inhibition observed with the parent compounds was likely not due to contamination with the oxon. Thus, OP bioactivation occurs at all planarian life-stages, including early during development/regeneration, similar to what is found in mammals (Timchalk et al. 2006). We did not observe significant differences in bioactivation of CPF and DZN between adult and regenerating planarians. Regenerating planarians retain DjChE activity in the ventral nerve cords throughout development, despite the loss of activity in the head. Normalized DjChE activity increases early during regeneration and plateaus around days 7-8 post-amputation. The results are in contrast with a previous study (De Simone et al. 1994) measuring ChE activity in the planarian species *Dugesia tigrina* where ChE activity in tail fragments was found to oscillate throughout regeneration roughly around the levels of intact worms. These differences may be due to differences in the experimental procedures or the planarian species used. Because of the lower total activity in regenerating planarians, we would expect them to be more sensitive to acute OP exposure than adult planarians, as has been found with rats (Timchalk et al. 2006). However, the increase in DjChE activity over the course of regeneration may be able to compensate for some of this during long-term exposure. It remains to be determined whether regenerating planarians can further modulate DjChE or downstream receptor expression as possible compensatory mechanisms in response to OP exposure, as has been seen in other models (Jett et al. 1993; Jameson et al. 2007).

### *D. japonica* planarians possess PON1 and CES activity

Using a standard biochemical assay, we demonstrated that *D. japonica* has calcium-dependent PON1 activity, which was comparable in magnitude to PTEH activity measured in mouse liver (42 pmol/(min*mg protein)) (Anspaugh and Roe 2002). Previous proteomic work has also suggested that a PON1 homolog exists in planarians (Geng et al. 2015). However, thus far, we have been unable to identify the encoding gene/transcript responsible for this activity in existing transcriptomic databases (Rozanski et al. 2019). Other proteins, such as squid Diisopropyl-fluorophosphatase (DFPase, EC 3.1.8.2), have also been shown to contain calcium-dependent PTEH activity (as measured here) and hydrolyze an array of organophosphorus nerve agents (Blum and Chen 2010), despite having low sequence similarity to PON1. Thus, it remains to be determined the exact identity of the planarian enzyme responsible for this activity, though the proteomic studies suggest some similarities with rat PON1, at least on a peptide level, exist.

We also characterized planarian CEH activity. We found minor inhibition in the CEH assay using the ChE specific inhibitor tacrine, which we have previously shown to inhibit DjChEs, albeit with lesser affinity than human AChE (Hagstrom et al. 2017), suggesting that DjChEs are not the primary contributor to the observed activity. In contrast, the CES specific inhibitor benzil (Hyatt et al. 2005; Wang et al. 2018) caused significant inhibition in the CEH assay, but negligible inhibition in the Ellman assay. Thus, DjChE and DjCES activity seem to be distinct. These findings agree with previous work in *D. tigrina* that showed enzymatic activity capable of hydrolyzing 1-naphthyl acetate that could be purified separately from ChE activity (De Simone et al. 1994). The finding that relatively high benzil concentrations (>3mM) caused only ∼40% inhibition in the CEH assay suggest that either *D. japonica* CES enzymes have structural differences in their binding sites compared to human CES, or other esterases may be partially responsible for the observed activity. To differentiate between these possibilities, future work needs to characterize the CES gene(s) using RNA interference, as we have previously done for DjChEs (Hagstrom et al. 2018).

### *D. japonica* has unique strengths as a non-animal HTS model for studying OP neurotoxicity

We have previously shown that *D. japonica* has two ChE genes with intermediate characteristics between AChE and butyrylcholinesterase (Hagstrom et al. 2017, 2018) that are sensitive to OP inhibition and can be reactivated by oximes. Furthermore, we have shown that different OPs have varying potency and cause different behavioral phenotypes in developing and adult planarians (Hagstrom et al. 2015; Ireland et al. 2022), suggesting that they affect a multitude of targets in addition to their shared effects on AChE, as seen in other systems (Dam et al. 2000; Slotkin 2006; Slotkin et al. 2006; Yang et al. 2008; Brown and Pearson 2015; Mamczarz et al. 2016; Schmitt et al. 2019). Here, we complement these studies and showed that *D. japonica* have the necessary enzymatic activity to bioactivate and detoxify OPs in adult and developing specimen and are sensitive to specific inhibitors (benzil, tacrine) known to inhibit mammalian CES and ChEs (Taylor and Radić 1994; Hyatt et al. 2005; Wang et al. 2018).

Compared to other popular non-animal organismal models, namely nematodes, fruit flies, and developing zebrafish, *D. japonica* has unique advantages as a model system for OP studies. For example, nematodes have low neural complexity with only 302 neurons (Friedman 2019), fruit flies have negligible PON1 activity (Healy et al. 1991), and developing zebrafish cannot bioactivate OPs before 3 days post fertilization (Yang et al. 2011). Thus, OP pharmacodynamics and multi-OP interactions in these alternative models may not be representative of human exposure. Together, these results support the use of *D. japonica* as a non-animal model for studying OP neurotoxicity and developmental neurotoxicity.

## Acknowledgments

We thank Veronica Bochenek for help with experiments, Dr. Alexander Baugh for use of his plate reader, and Kevin Bayingana and Yeh Seo Jung for help with planarian care. Research reported in this publication was supported by the National Institute of Environmental Health Sciences of the National Institutes of Health under Award Number R15ES031354 (to E.M.S.C). The content is solely the responsibility of the authors and does not necessarily represent the official views of the National Institutes of Health.

## Ethical standards

The manuscript does not contain clinical studies or patient data.

